# Tryptophanol, a novel auxin analog found in marine diatoms, enhances nitrogen assimilation

**DOI:** 10.1101/2025.06.24.661426

**Authors:** Dong-Sheng Zhao, Shengqin Wang, Yi-Cheng Xu, Yu-Ting Chen, Muhammad Ahsan Farooq, Sue Lin, Peng Cao, Junliang Li, Zhi-Wei Hu, Nan Li, Matteo Scarsini, Chuanling Si, Shu-Ming Li, Xiufeng Yan, Qiuying Pang, Chris Bowler, Hui-Xi Zou

## Abstract

Diatoms exhibit superior competitive capacity in nitrogen assimilation, largely contributing to their growth, although the mechanisms underpinning their success have not been completely understood. Here, a non-ribosomal peptide synthase-like (PtNRPS1, with an unusual domain structure A-T-R_1_-R_2_) gene was found to play a vital role in short-term nitrogen assimilation in the marine diatom *Phaeodactylum tricornutum*. *In vitro* biochemical assays and *in vivo* overexpression confirmed that PtNRPS1 catalyzed two sequential two-electron reductions of L-tryptophan to tryptophanol. Tryptophanol exhibits high structural and functional similarities to indole-3-acetic acid (IAA), the most typical phytohormone auxin. Surprisingly, the effective concentration of tryptophanol was lower than that of IAA by as much as 2-5 orders of magnitude for *P. tricornutum*. Compared with the action of IAA, a distinct molecular mode for tryptophanol was revealed by transcriptomic analysis, resulting mainly in enhanced short-term nitrogen assimilation, which was also confirmed by the elevated nitrogen uptake rates determined by stable-isotope tracking. Finally, global distribution of PtNRPS1 homologues from stramenopiles was found to be positively correlated with the abundance of genes involved in nitrogen assimilation pathways. Overall, our study provides evidence of an auxin-like derivative synthesized by an NRPS in a diatom. We speculate that tryptophanol may accelerate nitrogen assimilation, conferring advantages in the competition for nitrogen in the ocean.

## Introduction

Diatoms are photosynthetic members of the stramenopiles, which are of great ecological importance because they contribute to almost 20% of the world’s primary production.^1^ Their superior competitive ability for nitrogen, e.g., storing nitrate in excess of their immediate growth needs,^2, 3^ provides marine diatoms with certain advantages in their competition with other phytoplankton groups in the ocean environment. Moreover, diatoms tend to utilize nitrate for growth, while other phytoplankton groups, e.g., dinoflagellates, prasinophytes, and haptophytes, predominantly store it intracellularly.^4^ Yet, how diatoms assimilate nitrogen preferentially for their growth remains unclear.

Auxin was the first phytohormone identified in plants, and it plays crucial roles in plant growth and development.^5^ The chemical structures of auxins typically contain an indole ring, e.g., indole-3-acetic acid (IAA), which is the most abundant natural auxin. Besides terrestrial plants, auxin is also widely distributed in algae, including cyanobacteria, green algae, red algae, and brown algae.^6, 7, 8^ For diatoms, treatment with exogenous auxin or co-culturing with phycosphere-derived auxin-producing microorganisms can vastly promote their growth,^9^ suggesting the existence of a growth regulation mechanism by auxin in these phototrophs. However, to our knowledge a diatom-specific auxin has never been identified to date.^10^

Data mining of the increasing microbial genomic repositories has greatly accelerated the discovery of new natural products, e.g., terpenes, polyketides (PKs), and non-ribosomal peptides (NRPs).^11^ From the genetic perspective, the biosynthetic potential of natural products obviously far exceeds the number of compounds that have actually been isolated and identified.^12^ Using the data mining approach, it may be possible to discover novel secondary metabolites in diatoms, as the whole genome sequences of a growing number of marine diatoms have become available in the last few decades.^13, 14, 15^ Many of these diatoms possess NRP synthases (NRPSs) or NRPS-like enzymes, although the functions of these enzymes have not been investigated. The canonical NRPSs catalyze chemical reactions required for the activation and subsequent condensation of amino acid and/or carboxyl acid building blocks, normally using the adenylation (A) domains for substrate recognition, thiolation (T) domains to hold the activated substrate, and condensation (C) domains for peptide bond formation as the core domains to form the backbone of peptides, which would subsequently undergo modifications by optional domains, e.g., the methyltransferase, epimerase, and reductase domains.^16^

Previously, *in vivo* overexpression of *PtNRPS1* has enhanced tolerance of *Phaeodactylum tricornutum* to salicylate pollution, which provides this model marine diatom the potential for bioremediation.^17^ However, the biological function of PtNRPS1 is yet to be explored. In the current study, we demonstrate that PtNRPS1 is involved in the biosynthesis of tryptophanol in *P. tricornutum*. Because tryptophanol harbours an indole ring in its structure, we assessed the growth-promotion effect of tryptophanol and compared it with that of IAA. Subsequent analysis of the *P. tricornutum* transcriptome suggested that tryptophanol may enhance the growth of *P. tricornutum* through its role in nitrogen assimilation. Finally, the global distribution of PtNRPS1 homologues and the expression correlation between PtNRPS1 homologues and genes involved in nitrogen assimilation were analyzed using the *Tara* Oceans dataset. To the best of our knowledge, PtNRPS1 is the first example of a tetra-domain NRPS-like biosynthetic enzyme in eukaryotic algal secondary metabolism. Our results indicated the potential biosynthesis of NRPs in *P. tricornutum*, revealing the existence of tryptophanol in diatoms.

## Results

### PtNRPS1 plays important roles in short-term nitrogen assimilation

RNA-seq data from recent studies have shown that wild-type *P. tricornutum* cells can maintain a high level of *Phatr3_J47104* transcript in nitrate or nitrite media over a short time (15-45 min).^2, 3^ Interestingly, knocking out the nitrate reductase gene (*Phatr3_J54983*) can lead to impaired nitrogen assimilation from NO_3_^-^, resulting in the continuous expression of *Phatr3_J47104* at a high level in nitrate medium, as well as the accumulation of nitrate within the mutant cells over a long period (∼228 h).^2^ Based on these data from the literature, we speculated that the expression of *Phatr3_J47104* may correlate with nitrogen assimilation.

Quantitative analysis of *Phatr3_J47104* expression in *P. tricornutum* was performed to verify the results determined from RNA-seq data.^2, 3^ Compared with diatoms cultured in medium without an added nitrogen source, the expression of *Phatr3_J47104* in diatoms cultured in medium with added nitrogen (nitrite or nitrate) was upregulated over a short period (15 and 45 min) (Fig. 1a). In contrast, adding ammonium to the medium had little effect on the expression of *Phatr3_J47104*, although some increase in *Phatr3_J47104* expression was obvious at 45 min, and the increase was significant (Fig. 1a). Additionally, both ammonium and nitrite had little effect on *Phatr3_J47104* expression by 18 h, whereas nitrate still had some effect, as shown by the slightly upregulated expression by this time. Thus, these results suggest that *Phatr3_J47104* may play a potential role in nitrate assimilation, although which mechanism was involved remained unclear.

**Figure 1.**
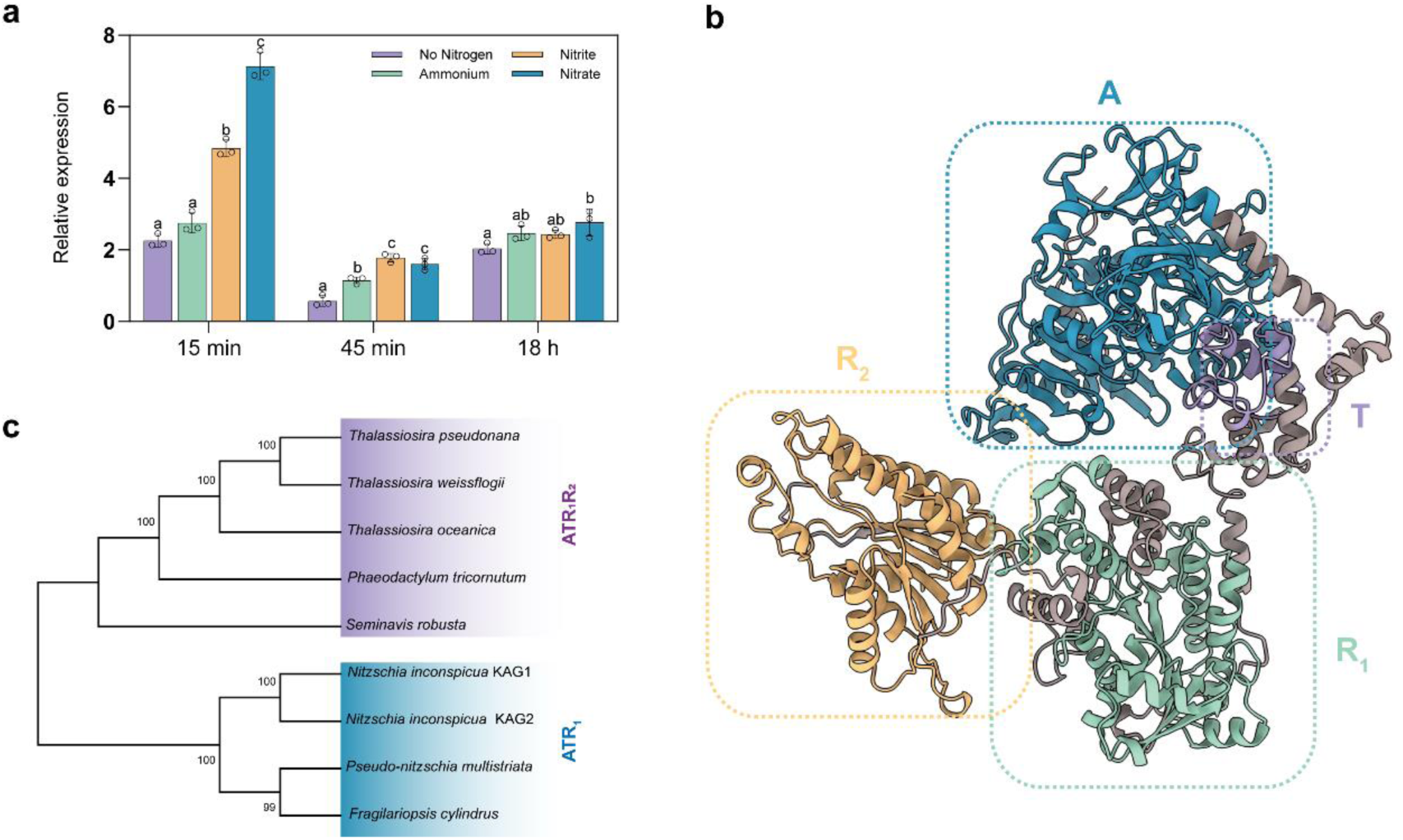
Gene expression and structure domains of PtNRPS1. **a** Transcriptional expression of PtNRPS1 under different nitrogen treatment conditions. Different letters denote significant difference between means (*p* < 0.05). Error bars indicate ±SE. **b** Domain architecture of PtNRPS1 harboring the four A-T-R_1_-R_2_ domains. A, adenylation; T, thiolation; R, reductase. **c** Phylogenetic analysis of PtNRPS1 and its homologues from genome-sequenced diatoms. Source data are provided as a Source Data file.

The Phatr3 annotation of the *P. tricornutum* genome indicates that *Phatr3_J47104* encodes a protein similar to non-ribosomal peptide synthases (NRPS),^13^ hence the name *P. tricornutum* (*Pt*) *NRPS1*. Conserved domain analysis revealed that PtNRPS1 was a single copy gene encoding four domains, one adenylation, one thiolation, and two reductase (R) domains (Fig. 1b and Supplementary Fig. 1). This unusual domain architecture (A-T-R_1_-R_2_) is widely spread in the fungi kingdom.^18^ Homologues can also be found in four other genome-sequenced diatoms besides *P. tricornutum*; *Thalassiosira pseudonana*,^19^ *Thalassiosira oceanica,*^20^ *Thalassiosira weissflogii*,^21^ and *Seminavis robusta*,^22^ suggesting the wide distribution of A-T-R_1_-R_2_ in diatoms (Fig. 1c and Supplementary Fig. 1). Additionally, homologues that lack the last R_2_ domain are encoded in the genomes of three other diatoms, including *Fragilariopsis cylindrus*,^15^ *Pseudo-nitzschia multistriata*^23^ and *Nitzschia inconspicua*^24^ (Fig. 1c and Supplementary Fig. 1). As NRPSs are usually involved in the biosynthesis of secondary metabolites, this raises the question of what products are synthesized by PtNRPS1 and its homologues in diatoms.

### PtNRPS1 prefers substrates with cyclic π bonds

The A domain of PtNRPS1 is expected to play a role in the activation and thioesterification of the correct substrate.^25^ Protein structure modeling of the A domain was used to assess potential substrate specificity. Specifically, AlphaFold V2.3.2 was used to build the model of the A domain of PtNRPS1, as well as that of AnATRR, the homologue of PtNRPS1 from the filamentous fungus *Aspergillus nidulans*.^18^ Domain A of both PtNRPS1 and AnATRR, together with 1AMU (Phenylalanine-activating subunit of gramicidin synthetase 1, PheA in *Brevibacillus brevis*),^26^ were subjected to structure comparison analysis (Supplementary Fig. 2). The A domain of PtNRPS1 was found to exhibit high structural and sequence similarity to the A domains that utilize aromatic amino acids as substrates (Supplementary Fig. 2). Aromatic amino acid residues such as Tyr70, Tyr84, Tyr91, and Phe215 of PtNRPS1, located near the substrate-binding site, could provide optimal positioning for aromatic rings, facilitating their proper alignment in the pre-reaction state, similar to that of PheA.^27^ These residues could also create favorable conditions for cation-π interactions with positively charged compounds.^18^ For instance, AnATRR displays preference for a small betaine as its substrate, which participates in cation-π interactions with three aromatic residues in the substrate pocket of the A domain.^18^ Comparative structural analysis of PtNRPS1, AnATRR and 1AMU revealed highly similar environments in their binding pockets (Supplementary Fig. 2). However, there are notable differences in the key residues, Trp204 and Phe202, that interact with betaine-like substrates. In AnATRR, Phe202 is involved in π-π stacking with Trp204, orienting both side chains towards the interior of the pocket, consequently reducing its volume. In contrast, Phe214 in PtNRPS1, which occupies the same position as Phe202 in AnATRR, does not participate in π-π stacking with Trp217 but instead stabilizes its side chain through atypical hydrogen bonds with surrounding residues such as Glu319 and Asn212, making it more accommodating to larger substrates. In addition, Trp217 in PtNRPS1 also tends to form π-π stacking with Phe285, resulting in a pocket structure akin to that of 1AMU.

To search for the real substrate of PtNRPS1, a library containing nearly 1,416 compounds was constructed, followed by virtual screening using AutoDock Vina (see Materials and methods and Supplementary Data 1). Six candidate substrates were obtained, including 2-hydroxy-5-nitrobenzoic acid, sulfosalicylic acid, 5-chlorosalicylic acid, tryptophan, phenylalanine, and tyrosine, which exhibited high binding energies and docking orientations close to the pre-reaction position. Among the six candidate substrates, tryptophan maintained the salt bridge and hydrogen bond with Glu319 because of its larger side chain, while simultaneously engaging in strong π-π interactions with both Tyr70 and Phe215 of PtNRPS1 (Fig. 2a). This could further shorten the distance between the carboxyl group of tryptophan and the high-energy phosphate groups of ATP to enhance binding and to enable the reaction. The binding modes of PtNRPS1_A with the other five substrates are shown and discussed in Supplementary Fig. 3.

**Figure 2.**
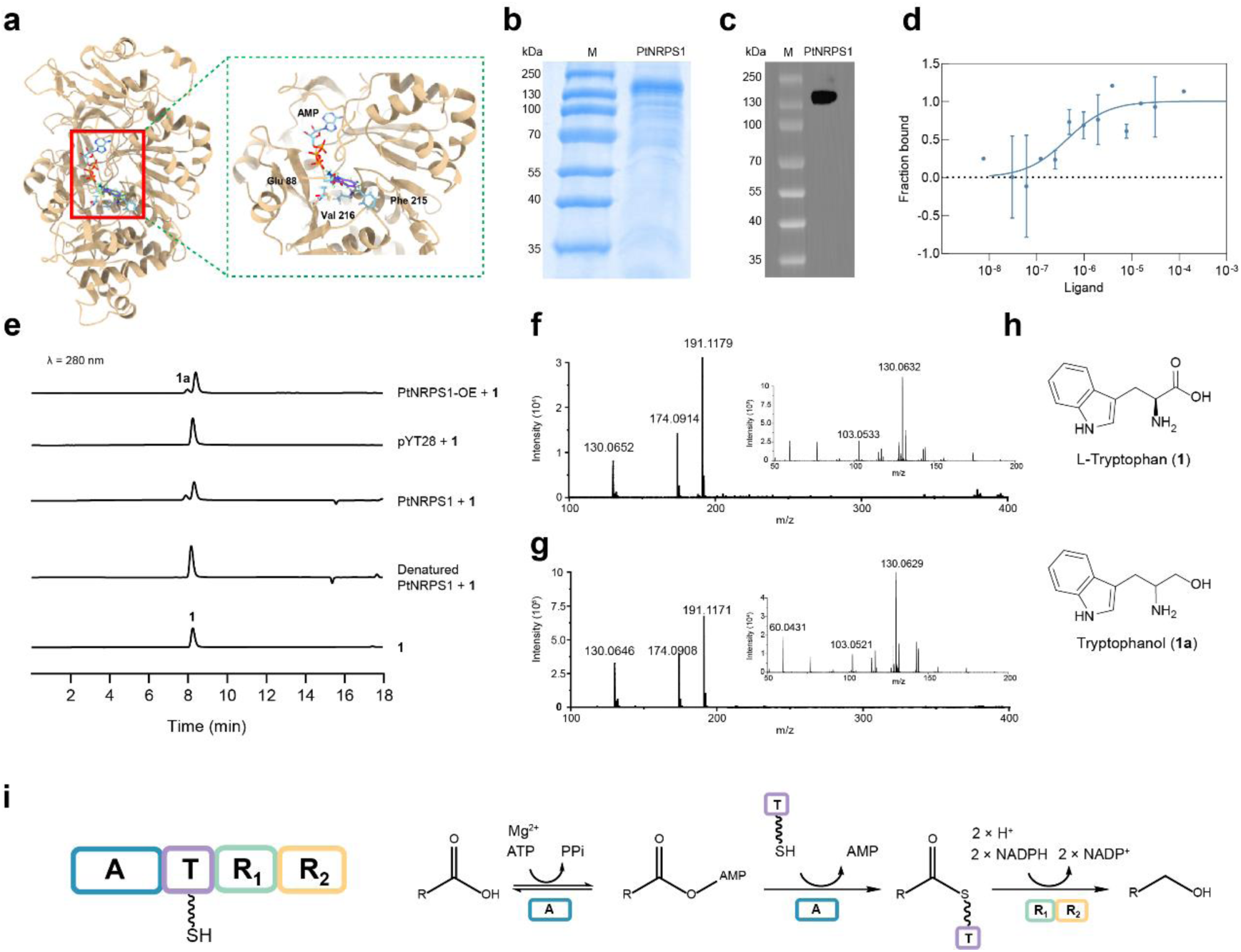
Biosynthesis of tryptophanol in *P. tricornutum* by PtNRPS1 using L-tryptophan as the substrate. **a** The structural details of PtNRPS1_A with L-tryptophan and AMP. Active site residues Glu88, Phe215 and Val216 directly involved in the positioning of L-tryptophan are shown as sticks. **b** SDS-PAGE analysis of the purified PtNRPS1 protein. **c** Western blot analysis of the purified PtNRPS1 protein. **d** Binding affinity curve of L-tryptophan with PtNRPS1 as determined by MST analysis. **e** HPLC analysis of the reaction mixture containing purified PtNRPS1 and L-tryptophan, and extracts of PtNRPS1-OE diatom cells fed with L-tryptophan, where denatured PtNRPS1 enzyme and *P. tricornutum* harboring pYT28 (without PtNRPS1 gene, see STAR Methods) was used as the control, respectively. **f** Mass spectra of the enzymatic product of PtNRPS1 incubated with L-tryptophan. **g** Mass spectra of the extra peak product of PtNRPS1-OE diatom cells fed with L-tryptophan. **h** Structures of L-tryptophan (**1**) and identified product (**1a**). **i** Domain architecture of PtNRPS1 and the postulated reaction. Source data are provided as a Source Data file.

Our previous work found salicylic acid (SA) and its 5-substituted derivatives (5-sSA) to be the best substrates for PtNRPS1 A domain (PtNRPS1_A) as determined by the hydroxamate assay,^17^ which does conform to the π-π interaction between PtNRPS1_A and the substrates. This work provided a rationale for us to use microscale thermophoresis (MST) to re-analyze the binding of PtNRPS1_A with the aromatic amino acids, i.e., L-tryptophan, L-phenylalanine, and L-tyrosine, all of which harbor a benzene ring with a cyclic π bond and are poor substrates for PtNRPS1_A according to the hydroxamate assay.^17^ As our initial attempts to get soluble recombinant PtNRPS1 protein and truncated forms of PtNRPS1 in *E. coli* were not successful, PtNRPS1 protein purified from PtNRPS1-OE diatom cells was instead used to perform the reaction (Fig. 2b and 2c). Surprisingly, the *K*_d_ value of the aromatic amino acid L-tryptophan was determined to be just 0.40 ± 0.29 μM (Fig. 2d), even lower than that of SA and 5-sSA.^17^

### Tryptophanol is produced by PtNRPS1 both *in vitro* and *in vivo*

Although SA and 5-sSA exhibited a high binding affinity for PtNRPS1, they are less likely to be the substrates of PtNRPS1, as confirmed in our previous study.^17^ To verify whether tryptophan, which exhibits higher binding affinity for PtNRPS1, could be accepted as a substrate by PtNRPS1, both *in vitro* enzyme reactions and *in vivo* transformations were carried out. When PtNRPS1 was incubated with L-tryptophan (**1**), a product peak (**1a**) was formed (Fig. 2e) and isolated for subsequent structural elucidation by MS and NMR (see Material and methods, Fig. 2f and Supplementary Fig. 4-7). Analysis by TOF-MS was carried out using both positive and negative ESI modes. In the positive mode, a [M+H]^-^ ion at *m*/*z* 191.1179 was obtained (C_11_H_14_N_2_O, -3.1 ppm, RDB = 6). According to the MS/MS spectrum from the high-resolution QTOF, the characteristic fragments were detected at m/z 130.0632 [M-C_2_H_6_NO]^+^, 103.0533 [M-C_2_H_6_NO-C_2_H_3_]^+^. The indole nucleus was further confirmed by the ^1^H-NMR spectrum, where the H-4 and H-7 protons appeared as doublets, H-5 and H-6 protons as triplets, and the noncoupled H-2 proton as a singlet (Supplementary Fig. 4). In comparison with that of L-tryptophan, the ^1^H-NMR spectrum of **1a** clearly showed the presence of signals for the extra methylene group at 3.61 and 3.44 ppm for the two H-12, indicating that the carbonyl group on C-12 of L-tryptophan was reduced. Finally, **1a** was identified as tryptophanol (Fig. 2h), and this was also confirmed by the ^13^C-NMR and 2D-NMR spectra (Supplementary Fig. 5-7).

The biocatalytic potential of PtNRPS1 *in vivo* was verified in PtNRPS1-overexpressing *P. tricornutum* cells constructed previously.^17^ An extra product peak was observed for L-tryptophan with a retention time of 7.9 min, which was identical to that of the *in vitro* enzyme product for PtNRPS1 incubated with L-tryptophan (Fig. 2e). The product was subsequently isolated for structural elucidation using TOF-MS (Fig. 2g). The spectrum obtained was identical to that of **1a**. The result therefore provided evidence of tryptophanol being produced as a product in PtNRPS1-OE cells when L-tryptophan was provided as a substrate. According to the experimental results, PtNRPS1 was able to convert a carboxylic acid to alcohol by two consecutive two-electron reductions (Fig. 2i).

### Tryptophanol promotes diatom growth and enhances short-term nitrogen assimilation

To assess the physiological effect of tryptophanol on *P. tricornutum*, bioassays were performed in which wild-type cells were treated with different concentrations of tryptophanol. Interestingly, the growth-enhancing effect of tryptophanol on *P. tricornutum* was extremely low as the concentration of tryptophanol increased (5 ∼ 100 pg L^-1^, Fig. 3a). However, at higher concentrations (≥ 1 ng L^-1^), tryptophanol significantly inhibited growth (Fig. 3a), suggesting the role of tryptophanol as an auxin analogue. To evaluate the growth-promoting effect of tryptophanol, the most important naturally occurring auxin, IAA, was chosen for comparison. In general, IAA can exert a growth promoting effect on *P. tricornutum* in the ranges from ng L^-1^ to mg L^-1^, with the minimum concentration being ∼ 1 ng L^-1,28, 29^ nearly 200 times higher than that of tryptophanol. Furthermore, IAA could induce the cells to reach a higher cell density, about 10% higher than the density of control cells after 96 h.^28, 29^ Although the maximum growth enhancement induced by tryptophanol was 6.45%, which is half that induced by IAA (10.66%), the concentration of tryptophanol yielding this enhancement was 4 × 10^-6^ times lower than that of IAA (20 pg L^-1^ *vs*. 5 μg L^-1^, Fig. 3a), suggesting that tryptophanol was a far more powerful growth-promoting agent than IAA.

**Figure 3.**
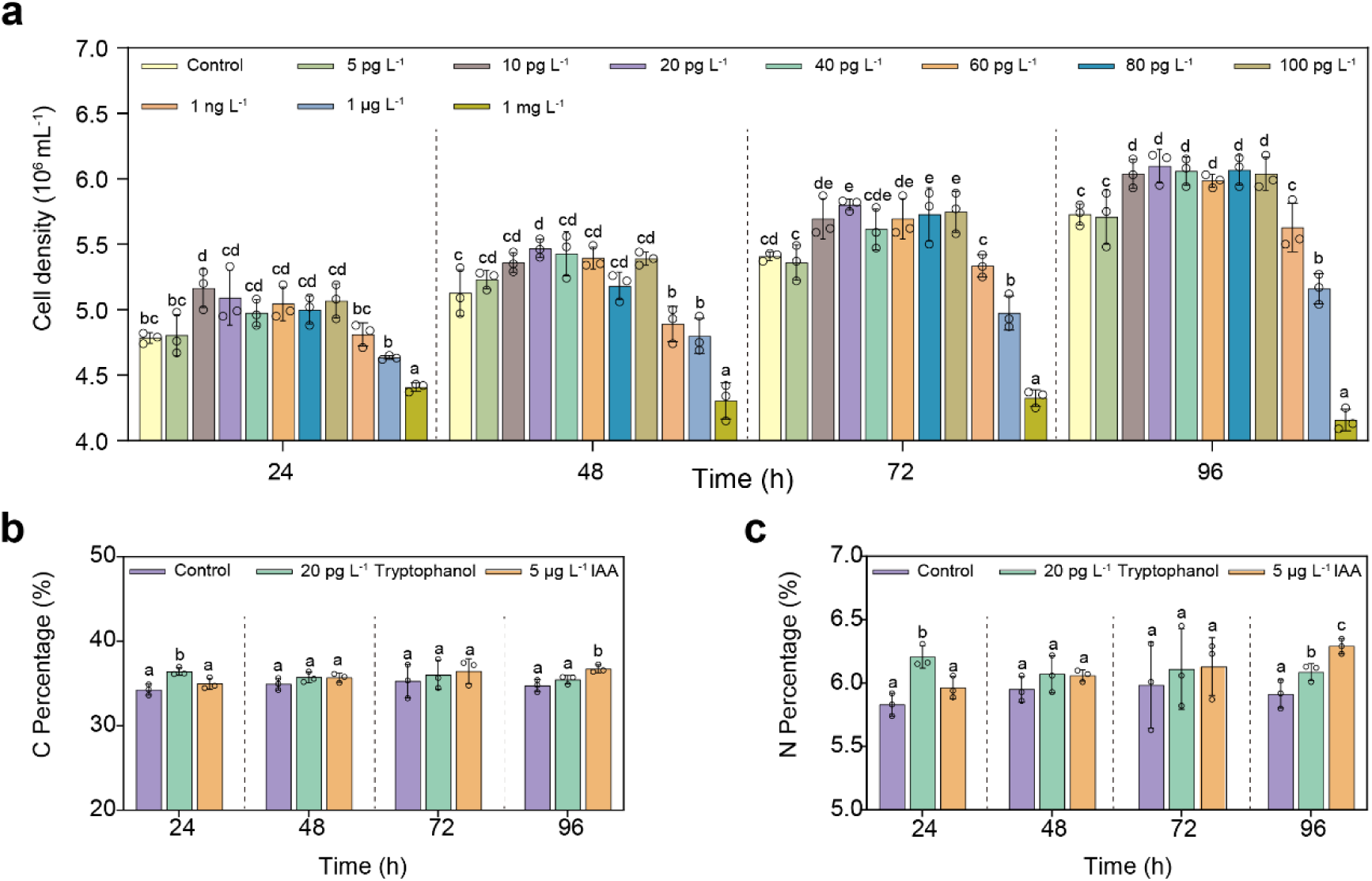
Growth promotion of *P. tricornutum* by tryptophanol and comparison of the effects of tryptophanol with that of IAA on organic matter content. **a** Changes in cell density of *P. tricornutum* cultures in response to treatments with different concentrations of tryptophanol. **b** Changes in total organic carbon contents in *P. tricornutum* cells treated with tryptophanol or IAA. **c** Changes in total nitrogen content in *P. tricornutum* cells treated with tryptophanol or IAA. Different letters denote significant differences between means (*p* < 0.05). Error bars indicate ±SE. Source data are provided as a Source Data file.

To examine whether organic matter in the cells could be affected by tryptophanol and IAA treatment, the content of total organic carbon (TOC) and total nitrogen (TN) was determined (Fig. 3b and c). Treatment with 20 pg L^-1^ of tryptophanol for 24 h significantly increased the content of organic matter, especially that of TN, which was 6.46 ± 0.92% higher than that of the control group. The content of TN was slightly increased by IAA treatment after 24, 48, and 72 h, but these increases as well as the increase observed for tryptophanol treatment after 48 and 72 h were not statistically significant. Tryptophanol also increased the content of TN after 96 h, but to a much lesser degree when compared with the increase after 24 h. Meanwhile, IAA dramatically increased the content of TN after 96 h, yielding a similar effect as that of tryptophanol after 24 h of treatment. These findings therefore indicated that tryptophanol could upregulate the pathway of nitrogen assimilation, leading to enhanced growth.

To obtain more insights into the mechanism underlying the growth-promoting effect of tryptophanol on *P. tricornutum* and to compare it with that of IAA, diatom cells were treated with tryptophanol and collected at the specific time points previously reported (every 24 h during the 4 day treatment),^29^ and subjected to transcriptomic analysis. Significant differences were found between the regulatory mechanisms of tryptophanol and that of IAA (Supplementary Fig. 8 and Supplementary Data 2).^29^ At 24 and 48 h, tryptophanol primarily affected metabolic pathways associated with ribosomes and oxidation, whereas IAA focused on carbon metabolism, secondary metabolite synthesis, cytosolic processes, and protein processing. At 72 h, tryptophanol primarily affected pathways such as the proteasome and carbon metabolism, while IAA primarily targeted secondary metabolism and oxidative processes. At 96 h, tryptophanol further influenced the synthesis of secondary metabolites and cofactors, while IAA treatment was associated with enhanced amino acid synthesis within carbon metabolism. In summary, tryptophanol promoted plant growth by modulating the expression of ribosome-related genes, while IAA accelerated plant growth through the regulation of carbon metabolism and the synthesis of primary metabolites.

Surprisingly, very few genes involved in nitrogen assimilation were found to be significantly upregulated by tryptophanol in the four tryptophanol-treated groups compared with that of the control group, which could not explain the higher level of TN in diatom cells after tryptophanol treatment (Fig. 3c). We speculated that the upregulation of genes involved in nitrogen assimilation may therefore occur at an earlier stage than the dramatic accumulation of nitrogen 24 h after tryptophanol treatment. Therefore, a short time-course (1 and 4 h) experiment was conducted. As expected, almost all genes involved in nitrogen assimilation were found to be significantly up-regulated (Fig. 4a). Transcriptomics and quantitative PCR validation confirmed the upregulation of a large number of genes, such as nitrate transporter (*Phatr3_EG02608*, *Phatr3_J26029*), nitrate reductase (*Phatr3_J54983*), nitrite transporter (*Phatr3_J13076*), nitrite reductase (*Phatr3_J38548*, *Phatr3_EG02286*), nitrite and sulfite reductase (*Phatr3_J12902*), and glutamine synthetase 2 (*Phatr3_J51092*) following tryptophanol treatment (Fig. 4a). All these results indicate that tryptophanol may act as a positive signal that affects the expression of genes involved in nitrogen assimilation, resulting in rapid nitrogen accumulation within diatom cells, which may then lead to the promotion of growth.

**Figure 4.**
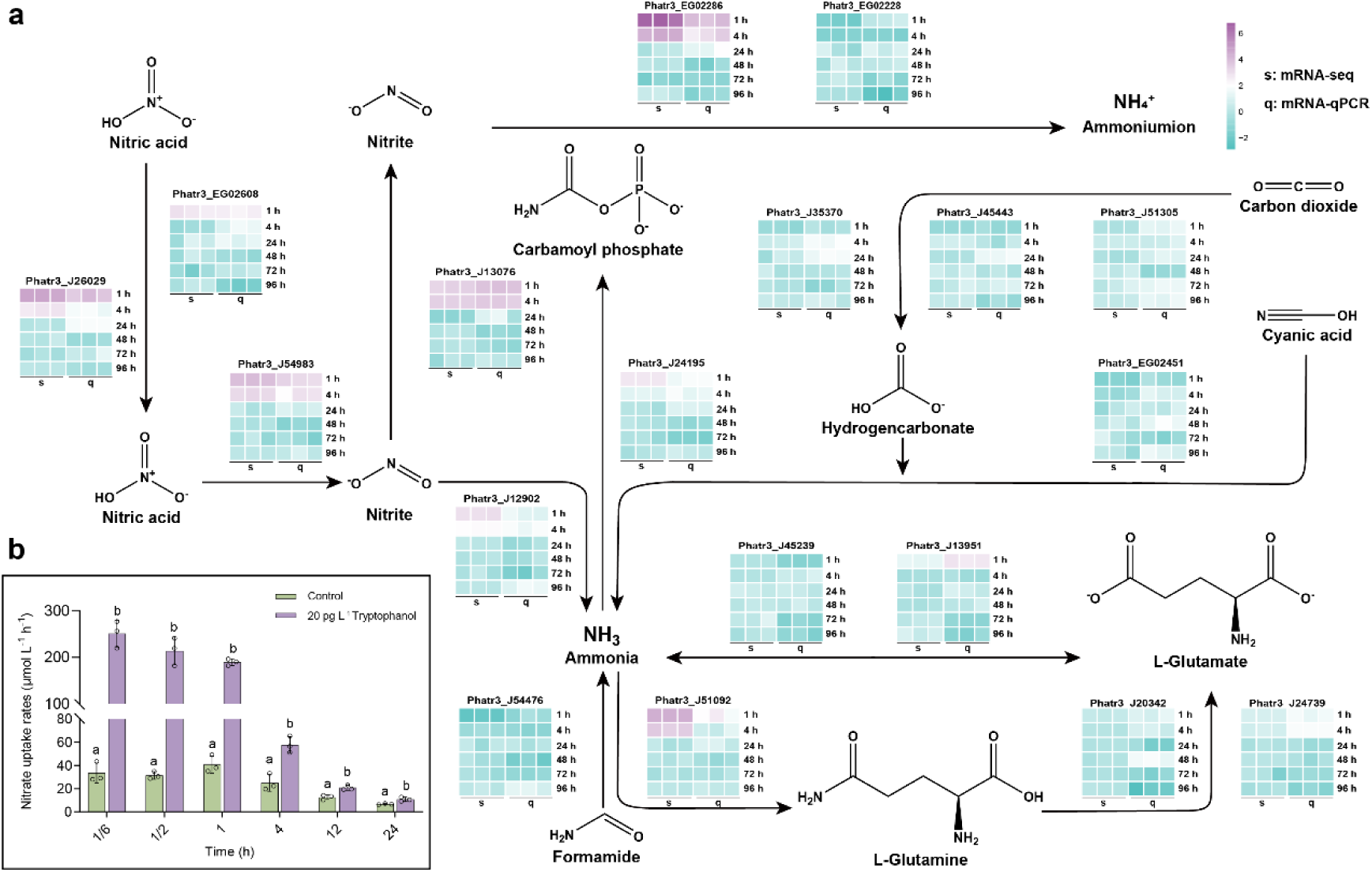
Effects of tryptophanol on nitrate uptake rates and expression of genes involved in the nitrogen assimilation pathway in *P. tricornutum*. **a** Transcriptomic analysis and qRT-PCR of genes involved in the nitrogen assimilation pathway of *P. tricornutum* under tryptophanol treatment. **b** Nitrate uptake rates of *P. tricornutum*. Different letters denote significant difference between means (*p* < 0.05). Error bars indicate ± SE. Source data are provided as a Source Data file.

To gather more evidence to support the ability of tryptophanol to enhance nitrogen assimilation in diatoms in the short term, the uptake rate of nitrate by *P. tricornutum* was further monitored using ^15^N-isotope labelled nitrate. The uptake rate was found to increase slightly within 1 h in the control (no tryptophanol) but then decreased slowly thereafter (Fig. 4b). On the other hand, treatment of *P. tricornutum* with 20 pg L^-1^ tryptophanol was found to rapidly enhance nitrogen assimilation in the diatom cells over a short duration, with the nitrate uptake rates being 7.92, 6.88, and 4.73 times greater than those observed in the control group at 1/6, 1/2, and 1 h, respectively (Fig. 4b). These results therefore confirm the role of tryptophanol in the short-term enhancement of nitrogen assimilation in *P. tricornutum*.

### Nitrogen assimilation pathway is positively correlated with PtNRPS1 homologues in stramenopiles in the global ocean

Not all PtNRPS1 homologues found in the sequenced genomes of diatoms contain two consecutive R domains, suggesting that these enzymes could not synthesize tryptophanol. To depict the distribution of PtNRPS1 homologues from stramenopiles in the global ocean, 23 sequences (see Supplementary Data 3), each of which possesses the four A-T-R_1_-R_2_ domains, were obtained and analyzed from *Tara* Oceans metatranscriptome dataset MATOUv1.5 and the eukaryote metagenomic dataset (Fig. 5a). PtNRPS1 homologues from stramenopiles were found in 269 of the total 581 samples (46.3%). Furthermore, through a separate analysis of *Tara* Oceans eukaryotic metagenome-assembled genomes (MAGs),^30^ we identified PtNRPS1 homologues in 14 MAGs from stramenopile (see Supplementary Data 3). This number may in fact may be larger due to the restricting filtering criteria to select only PtNRPS1 homologous sequences containing two consecutive R domains. Thus, our analysis does not exclude the possibility of the existence of PtNRPS1 homologues in other *Tara* Oceans samples. As *PtNRPS1* and its homologues are genes that are involved in the biosynthesis of secondary metabolites, they usually tend to be present in low abundance in the environment. This further suggests the wide distribution of PtNRPS1 homologues in the global ocean.

**Figure 5.**
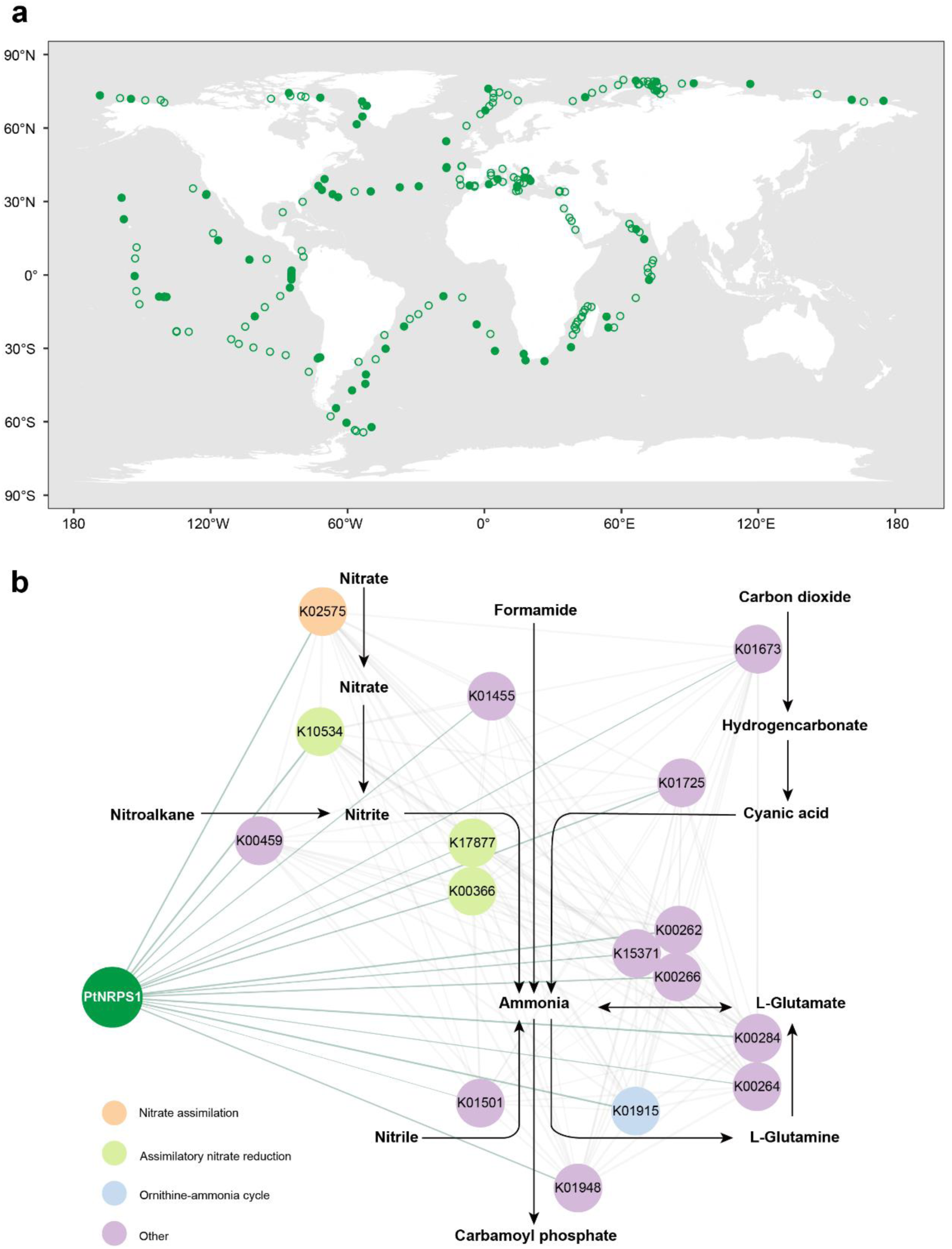
Global distribution of PtNRPS1 homologues in the *Tara* Oceans dataset and their co-expression network with nitrogen assimilation pathway genes. **a** Biogeographic distribution of Stramenopile-associated PtNRPS1 homologues in the *Tara* Oceans metatranscriptome MATOUv1.5. Solid and hollow circles indicate sampling stations with detected and undetected PtNRPS1 homologues, respectively. **b** Co-expression network between PtNRPS1 homologues and nitrogen assimilation-related KEGG Orthologs (KOs). Green lines denote PtNRPS1-KO correlations (rho ≥ 0.3, *p* < 4 × 10^-13^), and grey lines represent KO-KO correlations. Line thickness scales with correlation strength. Source data are provided as a Source Data file.

A total of 16 annotated KOs were identified as being involved in the nitrogen metabolism pathway in stramenopiles. Spearman correlation analysis was performed on the expression levels of these KOs and PtNRPS1 homologues (Fig. 5b). Consequently, expression of all the 16 KOs exhibited a significant positive correlation with that of PtNRPS1 homologues (rho ≥ 0.3, *p* value < 4 × 10^-13^). This could provide further evidence that PtNRPS1 homologues may play essential roles in nitrogen assimilation in stramenopiles, which may promote their growth in the ocean environment.

## Discussion

Tryptophan and its derivatives are common signaling molecules in the marine environment.^9, 31, 32^ In the present study, tryptophanol, a small organic molecule that is highly similar to tryptophan, was identified and its biological function in diatoms was characterized. Tryptophanol is known to be synthesized by the bacterium *Roseobacter* sp. via the transformation of diatom-derived organic matter, but the synthesized tryptophanol has only been identified by mass spectrometry without further structural elucidation.^33^ To the best of our knowledge, there has been no report documenting the isolation of tryptophanol from marine phytoplankton, especially diatoms. There is one study that suggests tryptophanol may be an important metabolite for alfalfa drought tolerance.^34^ However, the growth promoting effects of tryptophanol in plants or algae has never been reported.

IAA, the most abundant natural auxin, is also a derivative of tryptophan, and it was first isolated from marine organisms several decades ago.^35^ Although no direct evidence has been shown for the production of IAA by diatoms, it is usually produced by the microbes of the marine environment, and it can promote the growth of diatoms.^9, 28^ Interestingly, if the carbonyl group of tryptophanol is removed, the resulting structure and its growth-promoting effect will be highly similar to the structure of IAA. For terrestrial plants, the acidity in the apoplast is essential for the passive diffusion of IAA through the plasma membrane.^36^ However, IAA in the cytoplasm becomes charged and membrane-impermeable when the cytoplasm becomes more basic, and the transport of IAA out of the cell then requires auxin efflux carriers.^36^ Due to the chemical structure modification, tryptophanol is estimated to have a much higher p*K*_a_ than IAA, making it easier for tryptophanol to retain its protonation state. On this basis, tryptophanol is more likely to be transported back and forth across the membrane via passive diffusion. Additionally, we cannot exclude the possibility that diatoms have some specific transporters for tryptophanol, which need to be further investigated in the future.

Interestingly, not all diatoms are predicted to have the ability to produce tryptophanol because of the lack of the R_2_ domain in the PtNRPS1 homologues. It is worth noting that it is not uncommon for one microbe to exploit a useful substance that is produced by another microbe, as the exploiter microbe is unable to produce it.^37^ This may also be the scenario for tryptophanol-producing and -nonproducing diatoms.

Tryptophan is the main precursor for IAA (tryptophan-depend pathways), despite the existence of a tryptophan-independent pathway for the biosynthesis of IAA.^5^ Although not all genes nor the enzymes involved in these pathways have been fully characterized,^38^ the biosynthetic pathway of IAA is believed to involve multiple steps. Some homologous genes involved in the biosynthesis of IAA have also been identified in algae, though this does not necessarily mean that algae can produce IAA from tryptophan.^8^ In contrast, the one-step synthesis of tryptophanol by PtNRPS1 may give it great advantages in the biosynthesis of auxin-like compounds. From this perspective, tryptophanol may be a better choice than IAA with respect to growth promotion of marine diatoms, since tryptophanol-treated diatoms were also found to exhibit a greater capacity for nitrogen uptake in the short term.

The biocatalysis of carboxylate to alcohol usually involves two reduction steps.^39^ Carboxylate reduction is a challenging chemical transformation due to the thermodynamic stability of carboxylate. For instance, the general biosynthetic pathway of betaine from choline includes more than ten steps, which is high in energy costs, but the AnATRR-mediated reduction step can greatly shorten the pathway.^18^ Similarly, the cost of tryptophanol production by PtNRPS1 requires one tryptophan molecule and two NADPH molecules that provide reducing power. Hence, this one-step biosynthesis of tryptophanol dramatically helps marine diatoms to save energy by reducing the consumption of energy required for the synthesis of auxins necessary to promote growth. Within microalgae-bacteria consortia, bacteria utilize tryptophan released by microalgae to produce IAA, which feeds back to and induces the growth promotion of the host algae.^9^ Our results indicate that the presence of tryptophanol at picogram per liter concentrations was almost equally effective in terms of growth-promotion of diatoms achieved by IAA at the nanogram per liter concentration. Since neither IAA nor tryptophanol can be produced in precisely the same amount from the same amount of tryptophan, it is more cost-effective in terms of energy for diatoms to utilize tryptophanol rather than IAA for growth promotion. The extremely low effective concentration of tryptophanol may provide distinct advantages to phytoplankton in the marine environment, mainly by eliminating the consumption of nitrogen-containing metabolites, especially under nitrogen stress. For instance, the strategy adopted by *P. tricornutum* in response to thermal stress includes the activation of nitrogen metabolism,^40^ of relevance because ocean warming is often accompanied by a decrease in upper ocean nutrient availability.^41^

All the advantages cited above may make tryptophanol an ideal signaling molecule that effectively promotes diatom growth when the content of nitrogen instantaneously increases as a consequence of ocean upwelling, winter storms, river inputs, etc.,^42, 43, 44^ potentially leading to diatom blooms (Fig. 6). Although we have demonstrated the real possibility of the synthesis of tryptophanol from tryptophan by PtNRPS1 in a diatom, we have no direct evidence to indicate that tryptophanol is present in ocean water. Such evidence requires further work seeking to isolate the compound from the water. However, the effective concentration of tryptophanol that could trigger a growth-promoting effect for diatoms was extremely low, as our data indicated (Fig. 3a), which this might explain the failure to detect and isolate this compound from ocean water.

**Figure 6.**
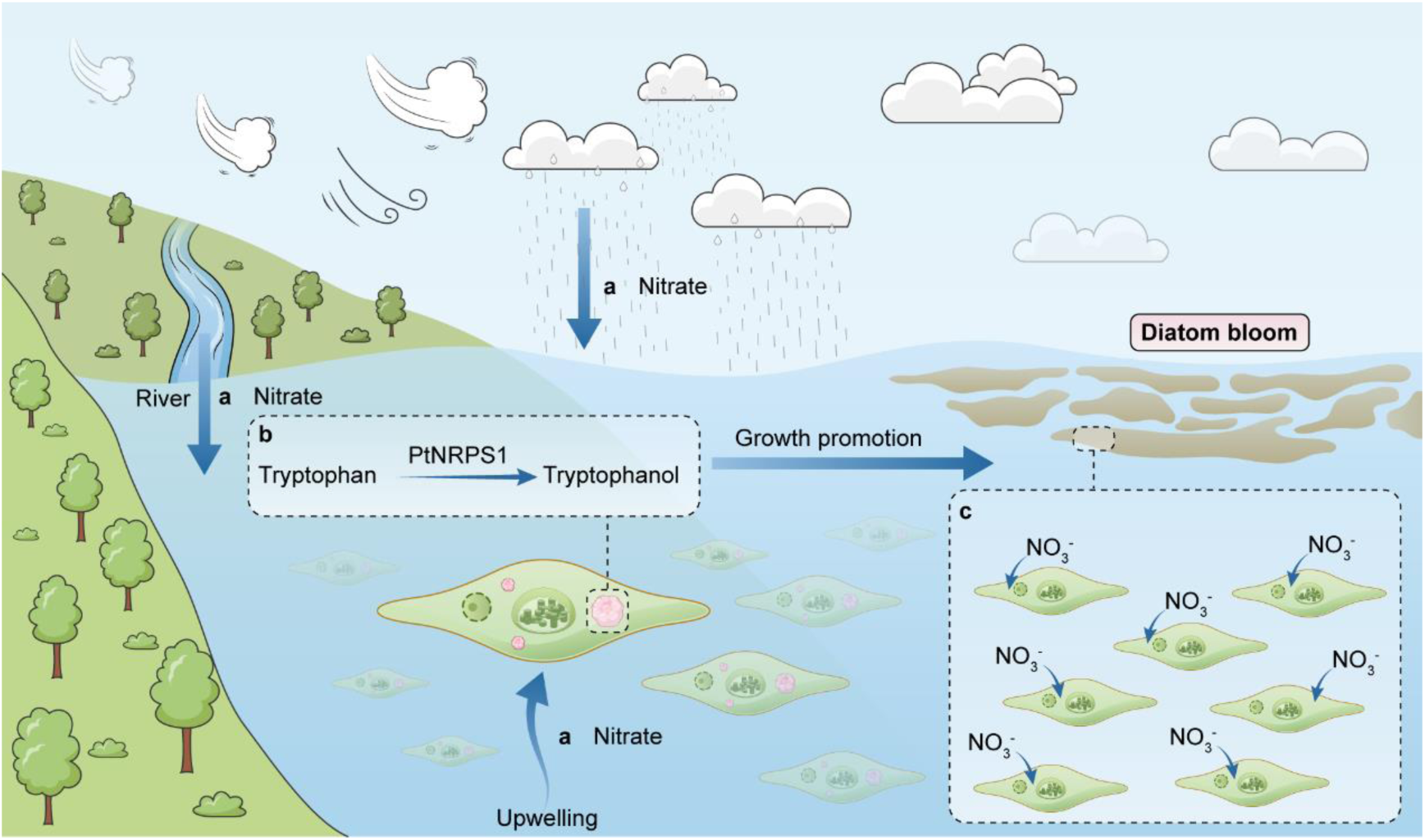
Proposed role of tryptophanol production on diatom blooms in the marine environment. **a** The expression of *PtNRPS1* homologues in diatoms is induced by periodic nitrate input in the ocean, largely from deep-ocean and coastal upwelling, river flow, dust storms, etc. **b** Tryptophanol is produced in diatoms possessing PtNRPS1 homologues. **c** Tryptophanol is released into the marine environment, which subsequently promotes the growth of the diatom community, potentially leading to diatom blooms.

In summary, we have discovered an unexplored non-ribosomal peptide synthase, PtNRPS1, in the marine diatom *P. tricornutum* and showed that it could catalyze two sequential two-electron reductions of L-tryptophan to tryptophanol. We also found that tryptophanol exhibited a significant growth-promoting effect, mainly via enhanced short-term nitrogen assimilation. Our work has for the first time provided evidence for an auxin-like derivative being synthesized by an NRPS in a diatom, with the effect of accelerating nitrogen assimilation to confer advantages in the competition for nitrogen acquisition in the ocean.

## Methods

### Chemicals and reagents

L-Tryptophan, IAA and ^15^N-labled sodium nitrate were purchased from Macklin (Shanghai, China). All other chemicals were obtained from Sigma-Aldrich (St. Louis, MO, USA).

### Microbial strains and culture conditions

The diatom *Phaeodactylum tricornutum* Bohlin CCMP2561 was used as the wildtype strain, which was obtained from the National Center for Marine Algae and Microbiota (USA). *P. tricornutum* that harbors pYT28*-*PtNRPS1 (PtNRPS1-OE) and *P. tricornutum* that harbors just the empty plasmid were prepared in our previous study.^17^ The diatom was grown at 20 ± 1 °C in f/2 medium but without Si. The medium contained the following components: 882 μM NaNO_3_, 36.2 μM NaH_2_PO_4_·H_2_O; Trace metal: 11.7 μM FeCl_3_·6H_2_O, 11.7 μM Na_2_EDTA·2H_2_O, 39.3 nM CuSO_4_·5H_2_O, 26 nM Na_2_MoO_4_·2H_2_O, 76.5 nM ZnSO_4_·7H_2_O, 42 nM CoCl_2_·6H_2_O, 910 nM MnCl_2_·4H_2_O; Vitamin: 296 nM thiamine·HCl, 2.05 nM biotin, and 0.369 nM cyanocobalamin. The diatoms were cultured under white, fluorescent lights (90 μmol m^-2^·s^-1^) with a 12 h:12 h dark-light cycle. For gene induction, phosphorous was omitted from the f/2 medium.

### RNA extraction and real-time quantitative PCR

Total RNA was extracted using Trizol (R0016, Beyotime Biotechnology, China) according to the manufacturer’s instructions. The extracted RNA was then used to synthesize the cDNA by reverse transcription using a HiScript III 1st Strand cDNA Synthesis Kit (Vazyme Bio, Nanjing, China). Quantitative real-time polymerase chain reaction (qRT-PCR) was conducted using the CFX96™ Real-Time System (Bio-Rad, Hercules, USA) and the HiScript II One Step qRT-PCR SYBR Green Kit (Vazyme Bio, Nanjing, China). The 18S ribosomal RNA gene was chosen as the internal control gene, and relative gene expression levels were determined using the 2^−ΔΔCt^ method.^45^ The sequences of the primers used in qRT-PCR are presented in Supplementary Table 1.

### Protein modeling for the PtNRPS1 adenylation domain and molecular docking

All models were built using AlphaFold (V2.3.2, https://colab.research.google.com/github/deepmind/alphafold/blob/main/notebooks/Al phaFold.ipynb),^46^ and visualized by Chimera X (https://www.cgl.ucsf.edu/chimerax/).^47^ The crystal structures of ligands were downloaded from PubChem (https://pubchem.ncbi.nlm.nih.gov/) in SDF format, which was then converted to Protein Data Bank (PDB) format using Open Babel program version 2.3.^48^ The geometrical parameters of enzymes and ligands were optimized without any constraint using GFN2-xTB method.^49^ Molecular docking was performed by the AutoDock GPU package,^50^ and the molecular graphics were prepared using Chimera X.

### Microscale thermophoresis analysis

Total protein was extracted from diatom cells using the BBproExtra^®^ Algal Protein Extraction Kit according to the manufacturer’s instructions (BestBio, Nanjing, China). The extracted proteins were used for microscale thermophoresis (MST) analysis performed on a Monolith NT.115 instrument (NanoTemper, Germany) as described previously.^17^

### PtNRPS1 activity assay

Total protein extracted from the diatom cells was concentrated by ammonium sulfate precipitation and dialyzed as described.^51^ PtNRPS1 was purified by FPLC using ÄKTA Purifier UPC10 (GE Healthcare, Pittsburgh, PA, USA) equipped with a HiLoad^®^ 16/600 Superdex^®^ 200 pg column (2.6 × 100 cm). The purity of PtNRPS1 was verified by SDS-PAGE and western blot and quantified using the Bradford method.^52^ The reaction mixture contained 50 mM Tris-HCl, pH 7.5, 10 mM ATP, 10 mM MgCl₂, 0.5 mM CoA, 1 mM NADPH, 10 mM substrate, and 0.5 mM PtNRPS1 protein. For the control sample, heat-inactivated PtNRPS1 was used instead. All reaction samples were incubated at 37 °C for 12 h, followed by three extractions with ethyl acetate (1:1, v/v). The organic phase was collected and centrifuged at 15,000 × *g* for 30 min to eliminate protein precipitates, and the resulting supernatants were pooled, evaporated to dryness, and reconstituted in methanol for HPLC analysis.

### Incubation experiments using *P. tricornutum* PtNRPS1-OE lines

Cultures (20 L) of *P. tricornutum* PtNRPS1-OE in the exponential phase (5 × 10^6^ cells mL^-1^) were collected by centrifugation at 4,000 × g for 5 min and then resuspended in the same volume of f/2 medium without phosphorous. Similarly, a control culture was also prepared; consisting of a *P. tricorntum* line carrying plasmid pYT-28. The feeding experiment was carried out by adding L-tryptophan to each culture to a final concentration of 10 mM. After 15 days of cultivation, the culture supernatant was extracted with ethyl acetate (1:1, v/v), which was repeated three times. The organic phase was collected from each extraction and centrifuged at 15,000 × g for 30 min to eliminate protein precipitate. The supernatants were then pooled, evaporated, and reconstituted in methanol for HPLC analysis and product isolation.

### HPLC conditions for analysis and isolation of products of PtNRPS1

HPLC was performed on an Agilent 1290 series system using an Agilent ZORBAX Solvent Saver High Definition (RRHD) StableBond-C18 column (150 × 3.0 mm, 1.8 μm) connected to a UV monitor set at 280 nm. The column was eluted with a system of solvents comprising double-distilled water (solvent A) and acetonitrile (ACN, solvent B), both of which contained 0.1% formic acid. A linear gradient of 5-100% (v/v) solvent B in 10 min and a flow rate of 0.4 mL min^-1^ were used. The column was then washed with 100% solvent B for 4 min and equilibrated with 5% (v/v) solvent B for 4 min between runs.

For product isolation, preparative HPLC was employed on an Agilent 1260 series system using an Agilent ZORBAX RX-C18 column (250 × 9.4 mm, 5 μm) with a flow rate of 4 mL min^-1^. The eluent was monitored by absorbance at 280 nm. Double-distilled water (solvent A) and ACN (solvent B), each containing 0.1% formic acid, were used as solvents. Elution was performed with a linear gradient of 5-60% (v/v) solvent B in 20 min. The column was then washed with 100% solvent B for 5 min and equilibrated with 5% (v/v) solvent B for 5 min in between runs.

### LC-MS/MS analysis of enzyme products of PtNRPS1

LC-MS was performed on a Sciex Exion system coupled with a Triple TOFTM 6600 mass spectrometer (Sciex, CA, USA). The LC consisted of a binary solvent system (solvent A: 0.1% formic acid in double distilled water; solvent B: 0.1% formic acid in ACN), an autosampler, and a column compartment. Chromatographic separation was performed on a Phenomenex Kinetex C18 reversed phase column (150 × 4.6 mm, 2.6 μm) at 40 °C using the following gradient: 0-1 min, 5% B; 1-10 min, 5-95% B; 10-12 min, 95% B; 12-12.1 min, 95-5% B; and 12.1-15 min, 5% B. The flow rate was set at 0.6 mL/min. The temperature of the autosampler was set at 15 °C and the injection volume at 5 μL.

The mass spectrometer was operated in positive ESI mode, and data were collected using information-dependent acquisition (IDA) featuring enhanced product ion (EPI) scan. Dynamic background subtraction (DBS) was also enabled. A TOF MS scan was performed at m/z 100-1200 Da. In each cycle, the 10 most intense ions over 50 cps were chosen for product ion scan at m/z 50-1200 Da. The optimized MS parameters were as follows: ion spray voltage of 5500 V; the turbo spray temperature of 550 °C; curtain gas of 35 psi; nebulizer gas of 55 psi; heater gas of 55 psi; declustering potential of 70 V; collision energy of 35 eV; and collision energy spread of 15 eV. In addition, an automated calibration delivery system (CDS) was used.

### NMR analysis and structural characterization of 1a

Samples used for NMR analysis were first dissolved in methanol-*d*_4_ or dimethyl sulfoxide (DMSO-*d*_6_) and subjected to Brucker AVANCE III 400 for taking NMR spectra. Both ^1^H- and ^13^C-NMR spectra were processed with MestReNova v11.0.4.

**1a** ^1^H NMR (400 MHz, Methanol-*d*_4_): δ 7.57 (1H, d, *J* = 7.9, H-4), 7.34 (1H, d, *J* = 8.1, H-7), 7.10 (1H, s, H-2), 7.09 (1H, t, *J* = 7.5, H-6), 7.01 (1H, t, *J* = 7.1, H-5), 3.61 (1H, dd, *J* = 10.1, 4.4, H-12), 3.44 (1H, dd, *J* = 10.8, 7.0, H-12), 3.20 (1H, m, H-11), 2.95 (1H, dd, *J* = 14.3, 6.2 , H-10), 2.76 (1H, dd, *J* = 14.2, 7.5 , H-10). ^13^C NMR (125 MHz, Methanol-*d*_4_): δ 138.24 (C-8), 128.83 (C-9), 124.29 (C-6), 122.43 (C-2), 119.72 (C-5), 119.33 (C-4), 112.31 (C-7), 112.02 (C-1), 66.53 (C-12), 54.49 (C-11), 29.68 (C-10) ppm. The NMR data of **1a** correspond well to those of tryptophanol.^53^

### Determination of the content of TOC and TN

Total organic carbon (TOC) and total nitrogen (TN) were determined as described,^54^ with some modifications. Briefly, diatom cells at the exponential phase (5 × 10^6^ cells mL^-1^) were collected by centrifugation at 4,000 × g for 5 min and then resuspended in the same volume of f/2 medium containing 20 pg L^-1^ tryptophanol or 5 µg L^-1^ IAA. A 50-mL sample of the culture was taken at 24, 48, 72, and 96 h, respectively, and diatom cells were collected by centrifugation at 4,000 × g for 20 min. After discarding the supernatant, the cell pellet was vacuum-dried. Then, 2 mg of the dried sample was weighed and subjected to TOC and TN content analysis using the Elementar Vario ISOTOPE elemental analyzer (Elementar, UK).

### Transcriptomic analysis

Total RNA integrity and concentration were determined by the RNA Nano 6000 Assay Kit on the Bioanalyzer 2100 system (Agilent Technologies, CA, USA) and the Nano-Photometer^®^ spectrophotometer (IMPLEN, CA, USA), respectively. RNA libraries were constructed with the TruSeq RNA Sample Prep Kit (Illumina, USA) and sequenced with the Illumina Hiseq 6000 platform. The reads with low and medium quality, with junctions, and those in which the base information could not be determined were filtered. The valid clean reads were mapped to the genomic contigs of the *P. tricornutum* genome (http://genome.jgi.doe.gov/Phatr2/Phatr2.download.html). The reference genome index was constructed using Hisat2 v2.0.5, and the paired-end clean reads were compared with the reference genome. Gene read counts were measured based on the Phatr2 (http://genome.jgi.doe.gov/cgi-bin/browserLoad?db=Phatr2) and Phatr3 (http://protists.ensembl.org/Phaeodactylum_tricornutum/Info/Index) gene models. Differential expression analysis (DEGs) was performed using DESeq2 R package (1.20.0), and the thresholds of |log_2_ (fold change)| ≥ 1 and *p* < 0.05 were used to identify significant DEGs. Genes were assigned to Gene Ontology (GO) databases and the Kyoto Encyclopedia of Genes and Genomes (KEGG) for functional and pathway analyses. Raw reads were submitted to the Sequence Read Archive database under accession numbers PRJNA1169328 (1 h and 4 h) and PRJNA1170410 (24 h, 48 h, 72 h, and 96 h). A description of sample IDs and accession numbers can be found in Supplementary Data 2.

### Nitrate uptake rate determinations

Exponential-phase diatom cells (5 × 10^6^ cells mL^-1^) were collected by centrifugation at 4,000 × g for 5 min and then resuspended in the same volume of f/2 medium containing ^15^N-labelled sodium nitrate as the nitrogen source. After adding 20 pg L^-1^ tryptophanol, 50 mL of the culture was taken at 1/6, 1/2, 1, 4, 12, and 24 h post-incubation and then centrifuged at 4,000 × g for 20 min. The cell pellet was washed with nitrogen-free seawater and again centrifugated at 4,000 × g for 20 min. The supernatant was then discarded, and the cell pellet was subjected to the same washing. The washing step was repeated three times. After the last wash, the cell pellet was vacuum-dried and then subjected to ^15^N absorption rate measurement using the Elementar Vario ISOTOPE elemental analyzer (Elementar, UK). The International Standards USGS-40 and USGS-41 were used to calibrate the instrument. Changes in nitrate levels in the culture medium and cells were determined using the commercial kits G0426W and G0409W (Grace Bio, Suzhou, China), respectively, according to the manufacturer’s instructions. The nitrogen uptake rate was calculated as previously described,^55, 56^ using the equation below:

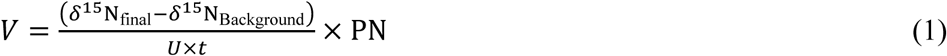

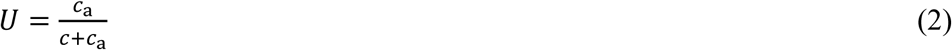

where *V* is the nitrogen uptake rate (μmol L^-1^ h^-1^), *δ*^15^N_final_ is the abundance of ^15^N in diatom cells of the treatment group, *δ*^15^N_final_ is the natural abundance of ^15^N (0.366%), PN is the concentration of nitrogen in diatom cells (μmol L^-1^), *U* is the relative proportion of nitrogen absorption (%), *c*_a_ is the concentration of ^15^N (μmol L^-1^), *c* is the concentration of dissolved nitrogen in medium (μmol L^-1^), and *t* is the cultivation time (h).

### Detection and correlation analysis of PtNRPS1 homologues in the *Tara* Oceans dataset

Accession numbers of the three types of domains (A, T, and R) collected from the Pfam library are shown in Supplementary Table 2. These domains were used as inputs to search for PtNRPS1 homologues in the Marine Atlas of *Tara* Oceans Unigenes dataset (MATOUv1.5, https://www.genoscope.cns.fr/tara/#MATOU-1.5),^57^ and *Tara* Oceans eukaryotic metagenome assembled genomes (SMAGs).^30^ After manual inspection, only hits harboring the intact four-domain (A-T-R_1_-R_2_) architecture, especially the two consecutive R domains, from stramenopiles were subjected to subsequent analysis. The distribution of PtNRPS1 homologues from stramenopiles in the global ocean was analyzed using MATOUv1.5.^58^ The abundance of PtNRPS1 homologues was calculated as the sum of the total gene coverages for each sample.

eggNOG-mapper was used for KEGG annotation of the UniGene obtained from MATOUv1.5.^59^ The normalized metatranscriptomic occurrences were also extracted from the MATOUv1.5 dataset.^57^ KEGG Orthology (KO) related to nitrogen metabolism was obtained from stramenopiles. The expression of PtNRPS1 homologues and each KO was calculated as the sum of the occurrences of all UniGenes involved in each group. The correlation between the expression of PtNRPS1 homologues and KO of nitrogen metabolism was then examined by Spearman’s rank correlation coefficient, whose co-expression network was constructed using Cytoscape.^60^ Additionally, samples were excluded from the correlation analysis where both the expression levels of the PtNRPS1 homologue and the KO group were zero.

### Statistical analysis

The experimental data were analyzed by OriginPro 9.0 and expressed as means ± standard deviations from triplicate (n = 3) determinations. Statistical significance was assessed with a two-way ANOVA followed by Tukey’s post hoc test to compare the results of the test group with those of the control group, and statistical significances were considered at the *p* < 0.05 and *p* < 0.01 levels. All statistical analyses were conducted using SPSS (Version 22.0).

## Data availability

All data supporting the findings of this study are available within the article, as well as the Supplementary Information file. Source data are provided in this paper.

## Author contributions

H.X.Z. and Q.P. conceived and designed the research; D.S.Z., Y.C.X, Y.T.C., S.L., P.C. and J.L. performed the experiments; Y.C.X., S.W., M.A.F., N.L., C.S. and Z.W.H. did the data analysis; C.B. and M.S. provided assistance with MATOUv1.5 and SMAGs analysis; H.X.Z. wrote the manuscript with contributions from all authors.

## Acknowledgments

This work was supported by Zhejiang Provincial Natural Science Foundation of China (LY23D060001), the National Natural Science Foundation of China (31670402), and the National Key R&D Program of China (2018YFD0901504). We thank Prof. Alan K. Chang (College of Life and Environmental Science, Wenzhou University) for his valuable discussion and for editing the manuscript, and Hai-Hong Wen (National and Local Joint Engineering Research Center of Ecological Treatment Technology for Urban Water Pollution, Wenzhou University) for assistance with experiments.

