## Supplemental information for "Tryptophanol, a novel auxin analog found in marine diatoms, enhances nitrogen assimilation"

Zhao *et al*.

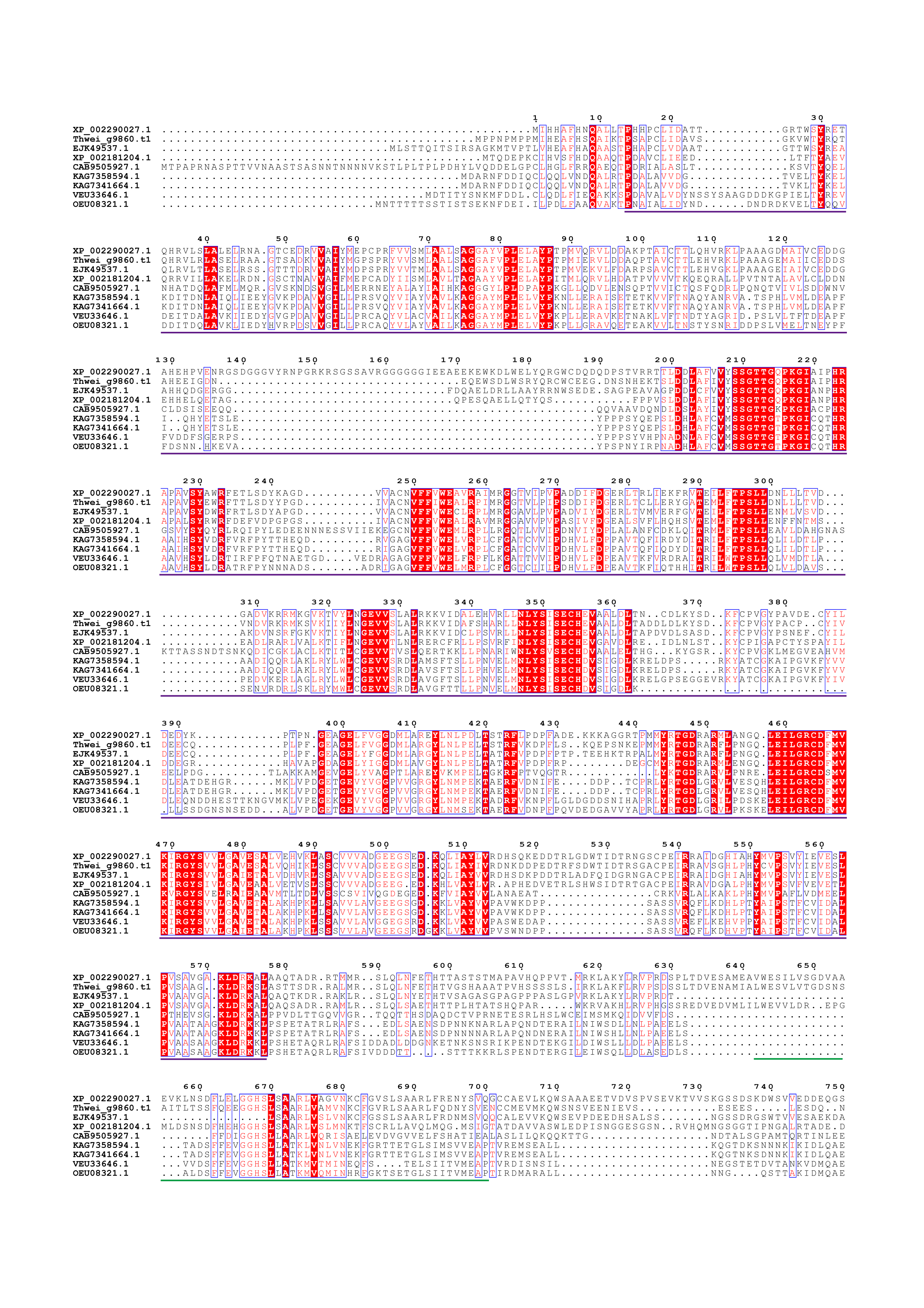

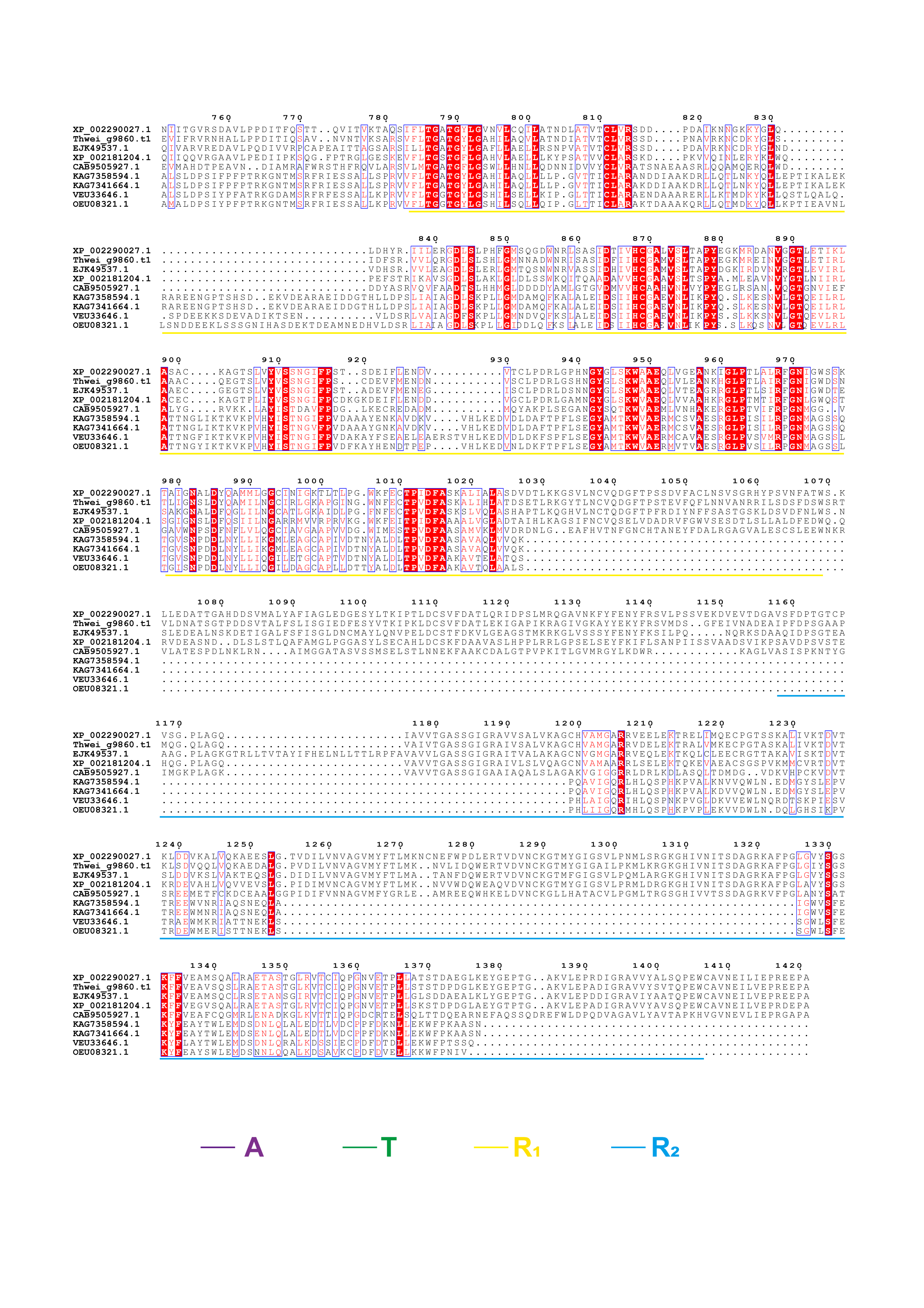

**Supplementary Figure 1. Sequence alignment of PtNRPS1 homologs in genome-sequenced diatoms.** *Thalassiosira pseudonana* XP_002290027.1, *Thalassiosira weissflogii* Thwei_g9860.t1, *Thalassiosira oceanica* EJK49537.1, *Phaeodactylum tricornutum* XP_002181204.1, *Seminavis robusta* CAB9505927.1, *Nitzschia inconspicua* KAG7358594.1, *Nitzschia inconspicua* KAG7341664.1, *Pseudo-nitzschia multistriata* VEU33646.1, *Fragilariopsis cylindrus* OEU08321.1. A, adenylation domain; T, thiolation domain; R, reductase domain. This figure is an extension to **Fig. 1c**.

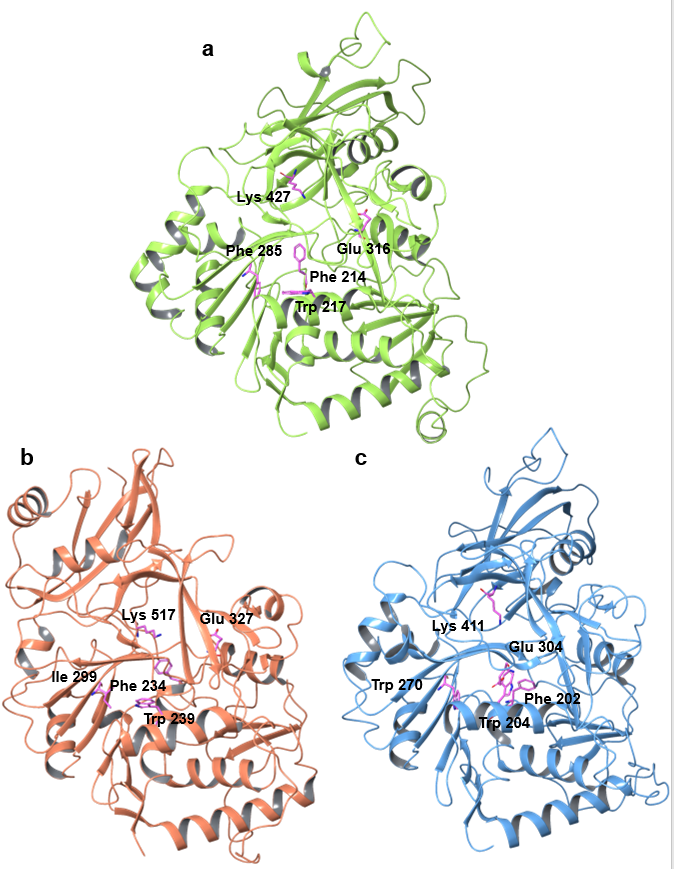

**Supplementary Figure 2. Structure comparison analysis of the AlphaFold2 models of the adenylation (A) domains of different NRPSs in complex with its cognate substrate L-phenylalanine. a** PtNRPS1 (from *Phaeodactylum tricornutum*) A domain. Residues facilitating the binding of tryptophan-like substrates through cation-π and electrostatic interactions are shown as sticks. **b** AnATRR (from *Aspergillus nidulans*) A domain. Residues facilitating the binding of betaine-like substrates *via* cation-π and electrostatic interactions are shown as sticks. **c** PheA (Phenylalanine-activating subunit of gramicidin synthetase 1, from *Brevibacillus brevis*) A domain (1AMU). Residues facilitating the binding of phenylalanine-like substrates through cation-π and electrostatic interactions are shown as sticks. This figure is an extension to **Fig. 2a**.

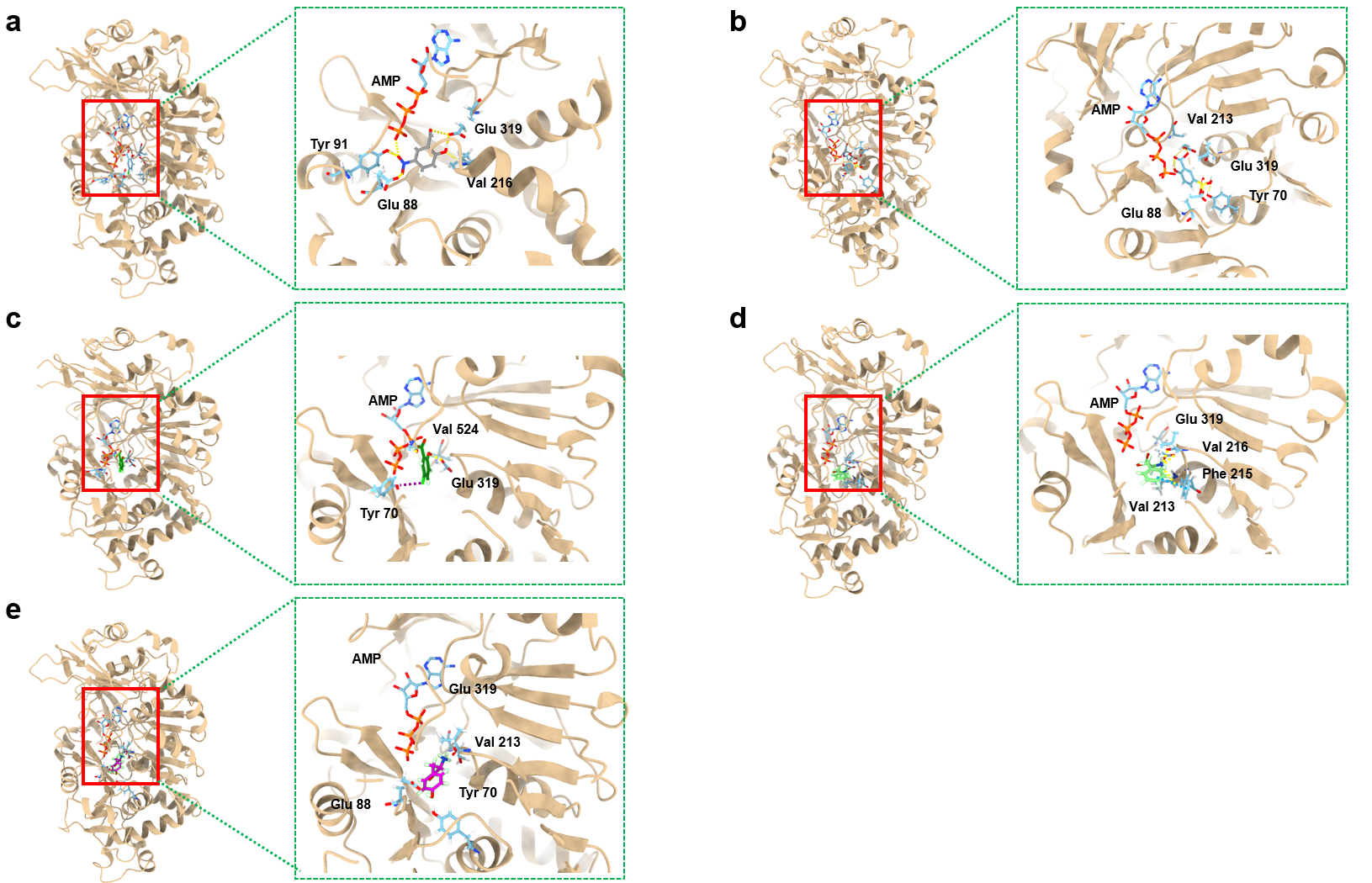

**Supplementary Figure 3. The structural details of PtNRPS1_A with the other five candidate substrates and AMP. a** 2-hydroxy-5-nitrobenzoic acid. **b** sulfosalicylic acid. **c** 5-chlorosalicylic acid. **d** phenylalanine. **e** tyrosine. In the process of phenylalanine binding to the A domain of PtNRPS1, Tyr70 and Phe215 play crucial roles in positioning the phenyl ring through π-π interaction. The carboxyl group of Glu319 forms both salt bridges and hydrogen bonds with the amino group of phenylalanine, bringing its carboxyl group into proximity with ATP, facilitating the reaction. Similarly, tyrosine binding is guided by π-π interactions with Phe215, positioning its aromatic ring. Additionally, its amino group forms salt bridges and hydrogen bonds with Glu319. However, unlike phenylalanine, tyrosine forms a hydrogen bond with Tyr70 via their hydroxyl groups, bringing tyrosine even closer to ATP, potentially increasing reaction efficiency. Sulfonyl salicylic acid exhibits a significantly different binding mode compared to the aforementioned aromatic amino acids. Its interaction with the adenylation domain primarily relies on a complex hydrogen bonding network involving the sulfonyl group and nearby residues, such as Tyr84, Glu88, and Val213. The docking results suggest that sulfonyl salicylic acid lacks specificity, potentially allowing it to bind to a broad range of NRPS adenylation domains and participate in reactions to a certain extent. The binding patterns of 2-hydroxy-5-nitrobenzoic acid and 5-fluorosalicylic acid show some resemblance to that of phenylalanine. The hydroxyl group adjacent to the carboxyl group mimics the amino group of amino acids, forming hydrogen bonds with Glu319 for positioning. However, due to the shorter distance between the carboxyl group and the aromatic ring, these compounds struggle to establish strong π-π interactions with Phe215. Instead, 2-hydroxy-5-nitrobenzoic acid relies on a nitro group to form a hydrogen bonding network with Glu88 and Tyr91, while 5-fluorosalicylic acid stabilizes its binding via a salt bridge between the fluorine atom and Tyr70. These data represent an extension to the results shown in **Fig. 2a**.

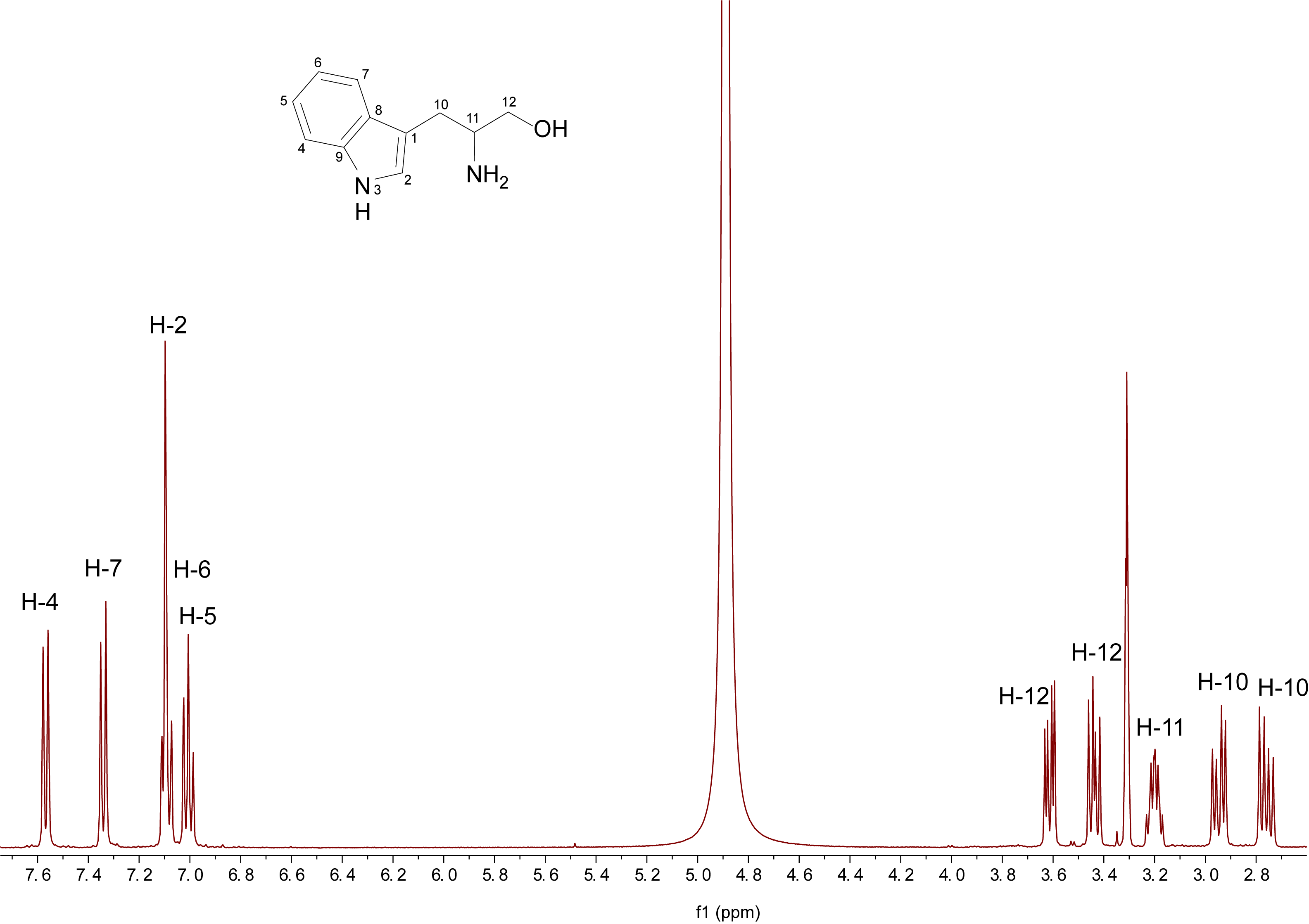

**Supplementary Figure 4. ^1^H-NMR spectrum of 1a in methanol-*d*_4_.**

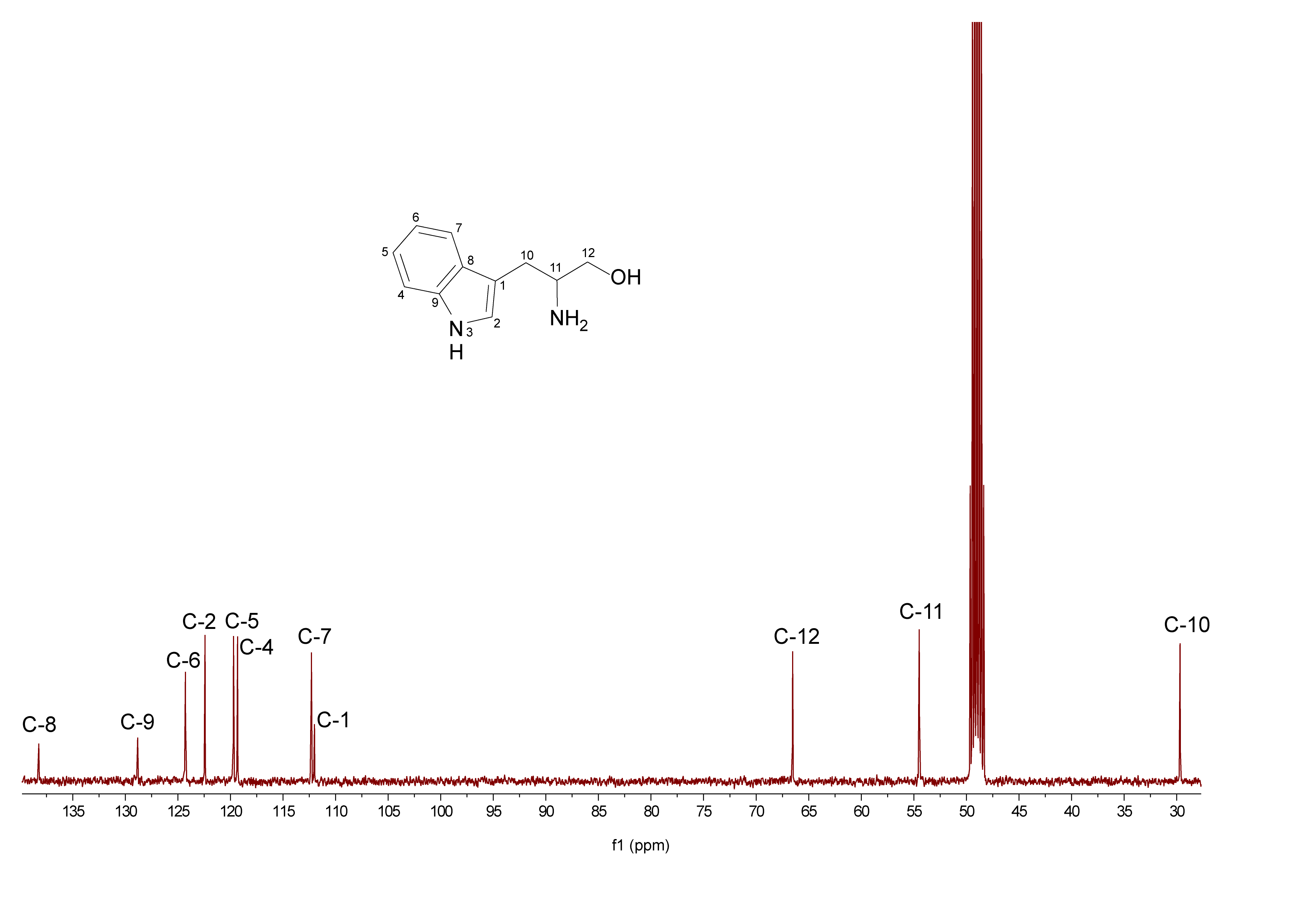

**Supplementary Figure 5. ^13^C-NMR spectrum of 1a in methanol-*d*_4_.**

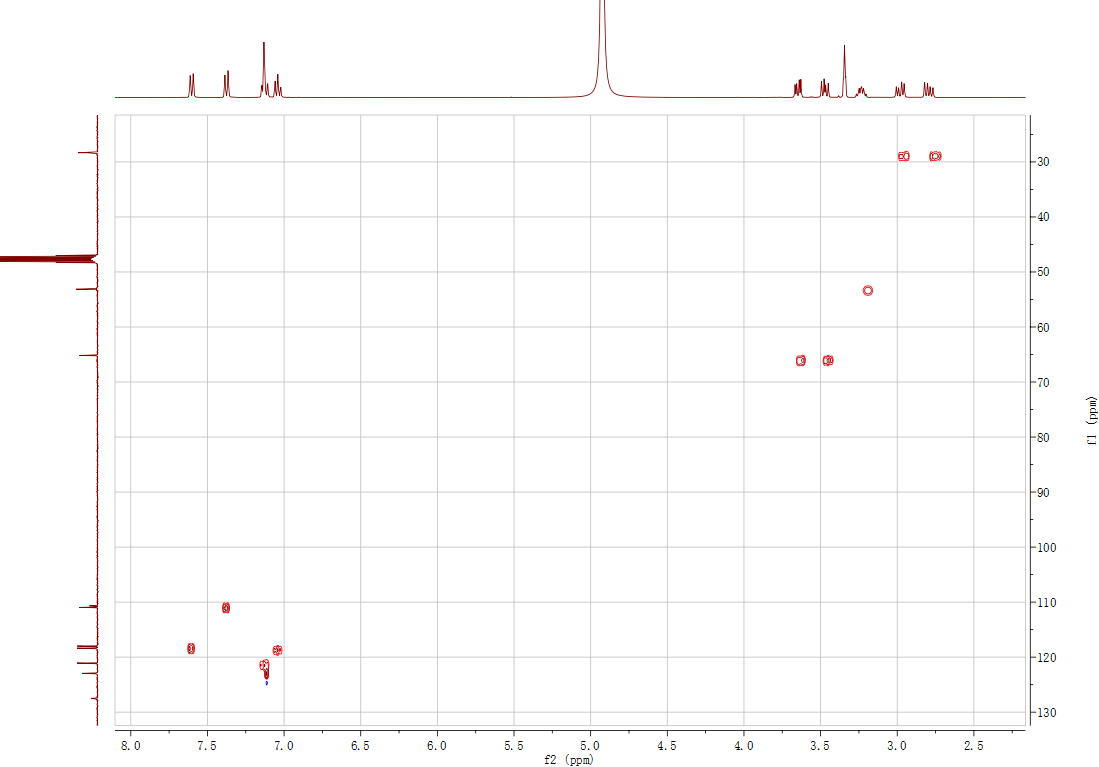

**Supplementary Figure 6. HSQC spectrum of 1a in methanol-*d*_4_.**

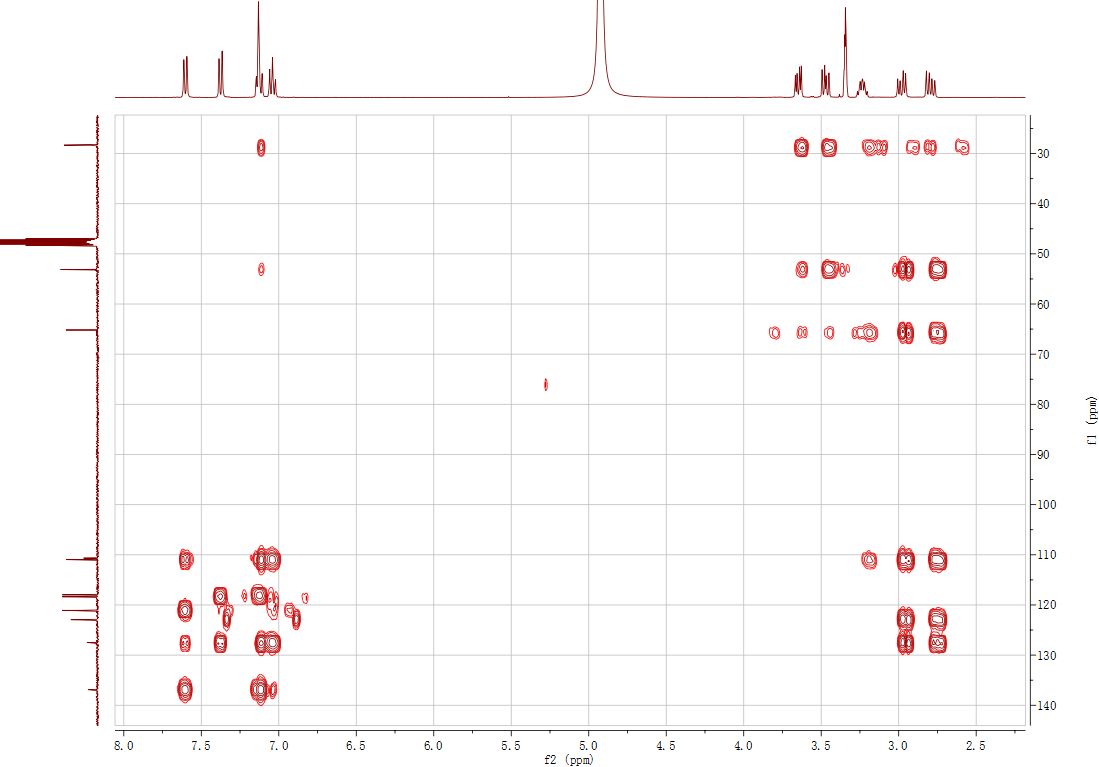

**Supplementary Figure 7. HMBC spectrum of 1a in methanol-*d*_4_.**

**Supplementary Figure 8. KEGG enrichment analysis of *P. tricornutum* treated with tryptophanol and IAA after 24 (a), 48 (b), 72 (c) and 96 h (d).** These data represent an extension to the results shown in **Fig. 4a.** Source data are provided as a Source Data file.

**Table S1. Primers used in this study.**

| Primer names | 5’ → 3’ |
| --- | --- |
| PtNRPS1_AT_for | gagctcggtacccctgacgccgtctg |
| PtNRPS1_AT_rev | caggtcgacaagctttacttctttgc |
| PtNRPS1_R_1__for | tggagctcggtacctttttgactggatcc |
| PtNRPS1_R_1__rev | gcaggtcgacaagcttcggctttatg |
| PtNRPS1_R_2__for | CCATCACGGATCCtctgcggtcgatccct |
| PtNRPS1_R_2__rev | AGATTAAGCTTccactcaggctgggaca |
| Phatr3_EG02608-F | CGCTTCTATTCCTACGGCATC |
| Phatr3_EG02608-R | AGATCCCATGACAAGCTGTG |
| Phatr3_J26029-F | GATAAGCCCACCCCAGTTG |
| Phatr3_J26029-R | AGCTACAAAGTCTGATCTCGC |
| Phatr3_J12902-F | GATCGAGAAGGCCAAGCTG |
| Phatr3_J12902-R | CAGCATACCGACGTACTTGAG |
| Phatr3_J54983-F | CTGAAGATCCCACAGTCAAGG |
| Phatr3_J54983-R | TCAAGAACGGCACAGGAATAG |
| Phatr3_J13076-F | TGCCCTTTGGTCTGATCATG |
| Phatr3_J13076-R | GGTTCCACTTTTGTATGCCAAG |
| Phatr3_EG02228-F | GACTGTTTGAATGTCGGATGC |
| Phatr3_EG02228-R | TGGTGACTGATTGTACTGCG |
| Phatr3_EG02286-F | ACGGTGTACAAGATGCAGTC |
| Phatr3_EG02286-R | GATGAGGCCGAAATCTTTGC |
| Phatr3_J54476-F | GTGCTCTGTCTACTTTCCTGTG |
| Phatr3_J54476-R | AAAAGAGACTTCACCATCCCC |
| Phatr3_J51092-F | TGATACCGTTCTTGCCGAATATG |
| Phatr3_J51092-R | TCGAAGAGCCATCAAAGTTCC |
| Phatr3_J24195-F | TGTTAACGTTTCCCCAGTAGG |
| Phatr3_J24195-R | ACCAGCACCTCCGAAATATAC |
| Phatr3_J45239-F | TTTTGACGAGGCAGGTGTAG |
| Phatr3_J45239-R | AGGACTTCGCTCACAATCTG |
| Phatr3_J13951-F | CGAGCGTATCATTGCCTTTC |
| Phatr3_J13951-R | GACTCCTTCATCGACAGTTGG |
| Phatr3_J20342-F | TTGGCGTTGTGTATGGGAG |
| Phatr3_J20342-R | AAAGCCAGTGTTAGTACGAGAG |
| Phatr3_J24739-F | GGTTTCCGACTACATTGCAAC |
| Phatr3_J24739-R | GAAGATCGTGGATAAGCTGGG |
| Phatr3_J45443-F | AATAAGTTCATTGCGGACAAGC |
| Phatr3_J45443-R | CTTCCGTGCCCATGATCAT |
| Phatr3_J51305-F | GAACAGTCCACTCTCCCAAG |
| Phatr3_J51305-R | ACATGACAGCCAAGTCGTTG |
| Phatr3_J35370-F | GACGGGACTTGAGGATGTTG |
| Phatr3_J35370-R | CAAGAGACAAATTCGCCACAC |
| Phatr3_EG02451-F | ATGGTATCATGAGTGCGATCG |
| Phatr3_EG02451-R | TCCGTTATCTTCCGCTTTCTG |

**Table S2. List of adenylation, thiolation and reductase related domains in Pfam library.**

| Accession number | Description | Domains |
| --- | --- | --- |
| PF00501 | AFD_class_I | A |
| PF13193 | AFD_class_I | A |
| PF14535 | AFD_class_I | A |
| PF01331 | Adenylation_DNA_ligase_like | A |
| PF16542 | Adenylation_DNA_ligase_like | A |
| PF01068 | Adenylation_DNA_ligase_like | A |
| PF09414 | Adenylation_DNA_ligase_like | A |
| PF00550 | PP-binding | T |
| PF07993 | NAD_binding_4 | R_1_ |
| PF01370 | Epimerase | R_1_ |
| PF01073 | 3Beta_HSD | R_1_ |
| PF04321 | RmlD_sub_bind | R_1_ |
| TPF16363 | GDP_Man_Dehyd | R_1_ |
| PF08659 | KR | R_2_ |
| PF00106 | adh_short | R_2_ |
| PF13561 | adh_short_C2 | R_2_ |

**Description of Supplementary Files**

**File Name: Supplementary Data 1**

**Description:** Ligand library for virtual screening for real substrate.

**File Name: Supplementary Data 2**

**Description:** Transcriptomic data.

**File Name: Supplementary Data 3**

**Description:** PtNRPS1 homologs possessing the four A-T-R_1_-R_2_ domains obtained from the *Tara* Oceans dataset.
